# Cortical Thinning Predicts Resting Vagally Mediated Heart Rate Variability: A Longitudinal Study Across Adolescent Development

**DOI:** 10.64898/2026.08.03.742432

**Authors:** Maximilian Schmaußer, Luise Baumeister-Lingens, Sonja Schulte, Michael Kaess, Romuald Brunner, Julian Koenig

**Affiliations:** University of Cologne, Faculty of Medicine and University Hospital Cologne, Department of Child and Adolescent Psychiatry, Psychosomatics and Psychotherapy, Cologne, Germany; University Hospital of Child and Adolescent Psychiatry and Psychotherapy, University of Bern, Bern, Switzerland; Department of Child and Adolescent Psychiatry, Centre for Psychosocial Medicine, University Hospital Heidelberg, Heidelberg, Germany; Clinic for Child and Adolescent Psychiatry, Psychosomatics and Psychotherapy, University of Regensburg, Regensburg

## Abstract

**Introduction:** Vagally mediated heart rate variability (vmHRV) reflects parasympathetic cardiac control and serves as a peripheral marker of brain-body interaction. While studies in adults link higher vmHRV to greater cortical thickness regions related autonomic regulatory, little is known about its association with longitudinal cortical maturation during puberty, a period of pronounced cortical thinning.

**Methods:** This longitudinal study examined whether individual differences in cortical thinning trajectories are associated with vmHRV in two independent cohorts of children and adolescents. Structural MRI were acquired in an accelerated longitudinal design over three time points, each one year apart in two cohorts (n = 44; ages 9 and 12 at baseline). Cortical thickness was estimated using FreeSurfer, and annualized regional thinning slopes were derived for 62 cortical regions. vmHRV was measured one year later at follow-up. Elastic net regression with stability selection identified robust predictors, which were entered into linear models separately for each cohort.

**Results:** Across both cohorts, vmHRV was associated with distributed patterns of cortical thinning. Consistent associations emerged in medial and posterior midline regions, including the precuneus, isthmus of the cingulate cortex, and medial prefrontal and orbitofrontal areas. Associations showed heterogeneous directions across regions, contrasting with uniform adult findings.

**Discussion:** vmHRV may be linked to network-level cortical maturation during adolescence, particularly within default mode and fronto-limbic systems. Findings extend adult work by demonstrating that brain-autonomic coupling emerges during development and is characterized by regionally differentiated trajectories of cortical thinning.

## Introduction

Heart rate variability (HRV) describes the natural fluctuations in heart activity from beat to beat (Berntson et al., 1997), with its high-frequency component, when measured at rest, representing a psychophysiological index of cardiac vagal activity (Kuo et al., 2005). The vagus nerve provides a structural and functional bidirectional connection between the brain and the heart (Thayer & Lane, 2009). Functional neuroimaging studies and human lesion studies have shown that multiple brain regions and networks are involved in the neural control of HRV (Thayer et al., 2012). Findings suggest a link between brain activity and HRV, as higher resting HRV is associated with greater functional connectivity between the amygdala and medial prefrontal cortex in younger and older adults (Sakaki et al., 2016). In the following, the term vagally-mediated HRV (vmHRV) is used to refer to high-frequency HRV at rest, which reflects parasympathetic cardiac control. Across the lifespan, HRV shows marked developmental changes (Finley et al., 1987; Finley & Nugent, 1995; O’Brien et al., 1986; Zhang, 2007), although there are relatively few longitudinal studies on normative HRV development in children and adolescents to date. Cross-sectional studies show that adolescents generally experience greater vmHRV than adults (Antelmi et al., 2004). It has been demonstrated that vmHRV increases during puberty until approximately age 18 before declining thereafter (Silvetti et al., 2001).

During adolescence, the brain undergoes fundamental restructuring, representing a crucial developmental phase for functional and structural maturation (Blakemore, 2012), characterized by “cortical thinning” (i.e., synaptic pruning; Koolschijn & Crone, 2013; Tamnes et al., 2010). Cortical thickness (CT) is considered to reflect cellular aspects of the cortical organization (Lerch, 2001) and has been suggested to be a sensitive method for detecting normative and pathological changes in brain structure, e.g. in contrast to measurements of brain volume (e.g., Thambisetty et al., 2010). The annual decrease in CT in individuals under the age of 20 is estimated to be over 1%, compared to 0.1% to 0.5% during the rest of life (Fjell et al., 2015). Previous work has generally observed a developmental decrease in CT from childhood (after age 2–3) to early adulthood (Walhovd et al., 2017), peaking at around age 8.5, independent of sex (Raznahan et al., 2011). Cortical regions implicated in the regulation of HRV in adults, such as the dorsolateral prefrontal cortex (PFC) and cingulate cortex, are the last to reach their maximum CT at approximately 10.5 years of age (Shaw et al., 2008). Longitudinal studies further indicate that rates of cortical thinning accelerate during adolescence (Ducharme et al., 2016; Tamnes et al., 2017). The process of cortical thinning is thought to reflect several underlying biological mechanisms (Jernigan et al., 2011), including increasing myelination of deeper cortical layers (Natu et al., 2019) and decreasing synaptic density (Huttenlocher, 1979).

Several studies have reported associations between CT in defined regions of interest (ROI) and vmHRV in adult subjects, implying associations of vmHRV with greater CT in cortical regions implicated in autonomic regulation, including the anterior cingulate cortex (ACC), orbitofrontal (OFC) and PFC (Winkelmann et al., 2017; Yoo et al., 2018). A large pooled mega-analysis further demonstrated parallel age-related declines in HRV and CT and showed that CT, particularly in the OFC, explained additional variance in HRV beyond the effects of aging (Koenig et al., 2021). Importantly, these associations were specific to HRV, as no significant relationships were observed between CT and heart rate (HR).

With regard to adolescence, a cross-sectional study by our group (Koenig et al., 2018) investigated age-related differences in the structural correlates of vmHRV in the brain. In a sample of female adolescents (n = 20, mean age 15.92 years) associations between vmHRV and CT in the right caudal ACC and rostral ACC (independent of hemisphere) were confirmed, resembling findings reported in adults. In contrast to adult samples, however, greater CT in all respective ROI was inversely associated with vmHRV, such that greater CT was related to lower vmHRV. These findings suggest developmental differences in the neural control of vmHRV and indicate a potential shift in the direction of the CT–vmHRV association across development (Koenig et al., 2018). Theoretically, greater vmHRV could be beneficial for cortical development during adolescence (i.e., cortical thinning) or could be partly result from pubertal cortical maturation processes (Carnevali et al., 2018).

Overall, higher vmHRV has been associated with greater CT in adults, whereas adolescence is characterized by pronounced cortical thinning alongside increases in vmHRV. Based on existing evidence on CT–vmHRV associations in adults (e.g., Winkelmann et al., 2017; Yoo et al., 2018) developmental cortical thinning during puberty does not appear to counteract the normative increase in vmHRV in adolescents (Koenig et al., 2021). While cross-sectional associations between CT and vmHRV have been reported, these findings cannot be transferred to the association between CT and HR (Koenig et al., 2018, 2021). Importantly, no study to date has longitudinally examined the development of CT (i.e., cortical thinning) and vmHRV across adolescence. Therefore, the present study aims to address this gap by investigating longitudinal trajectories of regional cortical thinning over three years of brain development and their associations with vmHRV. Given the lack of prior longitudinal evidence and region-specific hypotheses in adolescent samples, we adopted an exploratory, whole-brain approach to identify cortical regions in which developmental trajectories of cortical thinning are associated with vmHRV. Given the known sex differences in the development of cortical thickness (Koolschijn & Crone, 2013; Sowell et al., 2007) (Koolschijn & Crone, 2013; Sowell et al., 2007) and vmHRV (Koenig et al., 2017; Koenig & Thayer, 2016), we included sex as a confounding variable.

## Methods

The data for the present study were obtained from the Heidelberg Brain Maturation Study, an accelerated longitudinal cohort initiated in 2011 and conducted at the Department of Child and Adolescent Psychiatry at Heidelberg University Hospital in Germany. The study was approved by the Ethics Committee of the Medical Faculty of Heidelberg University (S-604/2011) and followed the guidelines of the Declaration of Helsinki (World Medical Association, 2013). In extension of the original study design, an additional evaluation was conducted in 2015 as a follow-up of the original sample, resulting in a total of four measurement time points. This follow-up study was also approved by the Ethics Committee of the Medical Faculty of Heidelberg University (S-046/2015). All participants and their legal guardians provided written informed consent prior to enrolment in the original study and the follow-up study. The participants received an allowance of €25 for the clinical assessment and €25 for the MRI session at each time point T1 to T3 and a total allowance of 75€ (telephone interview and study-sight visit) for the follow-up.

### Procedure

The original study comprised three visits to the study site at one-year intervals (T1-T3). These included cognitive assessments, magnetic resonance imaging (MRI), clinical interviews and psychological questionnaires. Assessments were conducted at the Department of Child and Adolescent Psychiatry, University Hospital Heidelberg, Germany. A clinical interview and a cognitive assessment were additionally conducted at the first appointment by trained psychologists. During this, participants and their parents completed various questionnaires on demographics, handedness, behaviour, temperament, and pubertal development (see Mürner-Lavanchy et al., 2020 for full reporting).

The follow-up examination (T4) included subjects who previously had at least two usable MRI data sets. The follow-up consisted of two parts, a telephone interview in which they completed a questionnaire. Subjects were then invited for a 60-minute onsite visit. They were first greeted and briefed before the ECG was set up for acclimatization. Subsequently, subjects completed computerized questionnaires and a physiological stress task. After removal of the ECG, each participant’s weight and height were measured.

### Participants

For the present study, two cohorts of participants aged 9 and 12 years were recruited. While the original study of Mürner-Lavachny et al. 2020 consisted of n = 112 individuals, n = 63 participants took part in the follow-up, of which n = 48 had complete and usable ECG data (Baumeister-Lingens et al., 2026). We report on a sample of n = 44 of which n = 21 form the older cohort (cohort 1) and n = 23 the younger cohort (cohort 2). Recruitment of participants took place on site, by mail, flyers or via the university’s website. The inclusion and exclusion criteria of participants were as follows: a) age of 9 or 12 years, b) right-handedness, c) German-speaking, d) psychiatric and neurological health (i.e. exclusion of all children with mental disorders), e) no psychotropic medication, f) birth weight > 2000 g, g) gestational age ≥ 36 weeks’ gestation, h) no mental disability (IQ ≥ 80) or developmental disorder (e.g. dyslexia), i) no braces, j) no twins and k) no siblings of previous participants.

### MRI Assessment

Whole-brain imaging data were acquired using a 3T Siemens Magnetom Biograph MRI scanner equipped with a 12-channel head coil at the Division of Radiology, German Cancer Research Center in Heidelberg, Anatomical T1-weighted images were obtained in the sagittal plane, comprising 192 slices with a 1-mm slice thickness and 1 × 1 mm² in-plane resolution (echo time [TE] = 2.52 ms, repetition time [TR] = 1,900 ms, flip angle = 9°). CT was assessed using automated cortical parcellation implemented in FreeSurfer version 5.3, employing the Desikan-Killiany atlas (Desikan et al., 2006). CT was initially estimated at each vertex as the shortest distance between the gray/white matter boundary and the gray matter/cerebrospinal fluid boundary on the reconstructed cortical surface (Fischl & Dale, 2000). Technical details of these procedures are provided in previous publications (Dale et al., 1999; Dale & Sereno, 1993; Fischl et al., 2001, 2002; Fischl, Salat, et al., 2004; Fischl, Sereno, & Dale, 1999; Fischl, Sereno, Tootell, et al., 1999; Fischl, van der Kouwe, et al., 2004; Han et al., 2006; Jovicich et al., 2006; Ségonne et al., 2004). Vertex-wise CT estimates were then averaged within each Desikan–Killiany region to derive regional CT measures used for statistical analyses.

All cortical reconstructions were visually inspected for segmentation accuracy, and datasets with major segmentation errors were excluded from further analyses.

### Physiological recording and data analysis

We report the recording and analysis of cardiac autonomic data in accordance with the “Guidelines for Reporting Articles on Psychiatry and Heart rate variability” (GRAPH; Quintana et al., 2016). The ECG was continuously recorded as a beat-to-beat interval (IBI) at 1024 Hz for the duration of the entire follow-up examination at T4 using an ECG-Move III chest strap (movisens, Karlsruhe, Germany). Data were visually inspected after each recording using the unisens viewer (version: 2.0 (Unisens - a Universal Data Format)) and stored in csv format. The ECG data were further processed in Kubios HRV 3.0 Premium (Tarvainen et al., 2014). A standardized 5-minute baseline segment was used for the further estimation of vmHRV. R-peak detection was manually corrected, and artifacts were removed. Smoothing priorities were selected as the detrending method (λ 500) for IBIs. The Kubios output was saved in txt format to later automatically read out the corrected IBIs and analyze HRV in R (García Martínez et al., 2017). IBIs corresponding to a mean HR < 30 or > 200 bpm were discarded. As a measure of vmHRV, the square root of the root mean squared difference of consecutive IBIs (RMSSD), a time domain measure, was determined (Task Force of the European Society of Cardiology the North American Society of Pacing Electrophysiology, 1996).

### Statistical Analyses

All statistical analyses were conducted in R (version 4.5.0) and performed separately for both cohorts. A region-based approach was employed, including all cortical regions defined by the Desikan–Killiany atlas. Regional CT (mean thickness within each ROI) was analysed separately for the left and right hemisphere. Cortical thinning refers to the developmental reduction in CT observed across adolescence. Analyses of regional CT included all 31captured regions, each for the left and the right hemisphere as depicted in Supplementary Table S1. Given the absence of a priori hypotheses regarding individual cortical regions and the unfavorable predictor-to-sample ratio, an exploratory, region-based analytical framework was adopted. To investigate the individual growth of regional CT over time from T1 to T3 of the n = 44 subjects and the relationship with vmHRV at T4, we used a multistep procedure designed to model individual change over time and apply regularized regression for variable selection. First, individual slopes for regional CT (in μm) were calculated for each participant using linear mixed-effects models across three time points (T1–T3), separately for each of the 31 brain regions, both hemispheres (left, right), and each timepoint, an MRI was taken (T1-T3). In these models, participants’ age at the three time points and participant ID were included as random effects, while sex was included as a fixed effect. This resulted in 62 predictors (2 hemispheres × 31 regions), representing subject-specific rates of cortical thinning per year. These were used as candidate predictors in all subsequent models. Given the limited sample size (n = 21 and n = 23), relative to the number of candidate predictors (62 regional cortical thinning measures), conventional multivariable regression would have been highly susceptible to multicollinearity, model overfitting, and unstable coefficient estimates. Therefore, rather than testing a large number of individual region-specific hypotheses, we adopted an exploratory, data-driven feature selection strategy that combined regularized regression with bootstrap stability selection. This approach was chosen to reduce model complexity, identify predictors that were consistently selected across repeated resampling, and generate robust candidate regions for subsequent inferential analyses. Accordingly, the present analyses are intended to be exploratory and hypothesis-generating in nature. Specifically, we applied elastic net regression with α = 0.5 and 10-fold cross-validation and a fixed random seed for reproducibility. Elastic net regression is a regularized regression approach that combines L1 (lasso) and L2 (ridge) penalties, thereby shrinking regression coefficients while accounting for multicollinearity among predictors, thus encouraging sparsity in the model and facilitating variable selection in high-dimensional data (Bühlmann & Geer, 2011; Zou & Hastie, 2005). The elastic net mixing parameter (α) determines the balance between ridge (α = 0), which preferentially shrinks correlated predictors together, and lasso regularization (α = 1), which promotes sparse variable selection by shrinking some coefficients exactly to zero. An intermediate value of α = 0.5 was chosen to balance predictor selection with coefficient stabilization in the presence of correlated cortical thickness measures. All regional thinning slopes were entered as predictors, and vmHRV was entered as the outcome variable. The regularization parameter (λ) was determined using 10-fold cross-validation. Predictor selection during bootstrap stability analyses was based on the more parsimonious λ corresponding to the one-standard-error rule. To assess the stability of predictor selection, the elastic net procedure was repeated 2,000 times using bootstrap resampling with replacement. In each iteration, the model was refitted using the λ corresponding to the one-standard-error rule, and selected predictors were identified based on non-zero coefficients. Predictors were considered stable if they were selected in a sufficient proportion of models (i.e., ≥ 20% of bootstrap iterations). As an additional sensitivity analysis, we evaluated the robustness of the selected cortical regions with respect to the choice of the elastic net mixing parameter. The complete bootstrap stability procedure was repeated across α values ranging from 0.1 to 0.9 (increments of 0.1), while all other model parameters, including λ selection, bootstrap resampling, and stability thresholds, remained unchanged. Selection frequencies were compared across α values to determine whether the identified cortical regions were consistently selected irrespective of the balance between ridge and lasso penalization. Results of these sensitivity analyses are reported in Supplementary Tables 6 & 7.The stable predictors were then entered into an ordinary least squares (OLS) regression model. Given high numbers of detected predictor regions, we performed automated model selection based on information-theoretic criteria using the dredge() function from the *MuMIn* package to identify the most parsimonious model for each cohort. Retained predictors were assessed for statistical significance (p < .05). To quantify the relative contribution of each significant region to explained variance in vmHRV, we calculated relative importance metrics using the lmg method from the *relaimpo* package. The *lmg* metric reflects the average increase in R² attributable to a predictor across all orderings in hierarchical regression models (Lindeman, Merenda, & Gold, 1980). To control for potential confounding effects, we additionally extended the original models by including the individuals’ age at time of HRV assessment (T4), the individual age-related slope of total brain volume, and sex as an interaction term with the selected regional predictors. Individual brain volume slopes were derived analogously to regional cortical thinning slopes, using linear mixed-effects models across T1– T3 with random effects for participant and age. Statistical reporting follows the APA Task Force on Statistical Inference guidelines (Wilkinson, 1999), ensuring transparency in reporting model selection procedures, effect sizes, and overall model performance.

## Results

At T1, cohort 1 (*n* = 21; 47.6% female) had a mean age of 12.58 years (SD = 0.34), whereas cohort 2 (*n* = 23; 43.5% female) had a mean age of 9.68 years (SD = 0.31). At T4, mean RMSSD was 53.97 ms (SD = 37.60) in cohort 1 and 54.54 ms (SD = 27.85) in cohort 2. Mean cortical thinning rates per year of the investigated brain regions are reported in Supplementary Tables S2 and S3. Using individualized cortical thinning slopes across 31 bilateral regions, we identified distinct spatial patterns of regional CT associated with interindividual differences in resting-state vmHRV.

### Cohort 1: development from 9 to 11 years

In the older cohort (cohort 1), vmHRV was significantly associated with cortical thinning in a distributed network of frontal, parietal and cingulate midline structures (see *Figure 1*). Specifically, greater cortical thinning in the left precuneus, *β* = −24.18, *SE* = 5.53, *t*(14) = −4.37, *p* < .001, was linked to higher vmHRV. In the isthmus of the cingulate cortex, *β* = −13.20, *SE* = 5.83, *t*(14) = −2.27, *p* = .038, as well as in rostral portions of the right anterior cingulate, *β* = −18.13, *SE* = 5.97, *t*(14) = −3.04, *p* = .007, cortical thinning showed consistent associations with higher vmHRV. In the left medial OFC, cortical thinning showed opposing associations with greater cortical thinning predicting lower vmHRV, *β* = 12.88, *SE* = 5.63, *t*(14) = 2.29, *p* = .03. The final model accounted for 65% of the observed variance in vmHRV (see Table 1). Sensitivity analyses demonstrated that the principal cortical regions identified by the stability-selection procedure were largely robust across a broad range of elastic net mixing parameters (α = 0.1–0.9), supporting the stability of the observed regional pattern (Supplementary Table 6). Including additional covariates did reveal significant effects of age at HRV assessment and global brain volume change over time whereas sex had no significant effect on HRV. However, the extended model did not provide a better fit compared to the original model, *F*(9, 4) = 0.72, *p* = .69, indicating that the inclusion of these variables did not improve explanatory power. For detailed results, see Supplementary Table S4.

**Figure 1.**
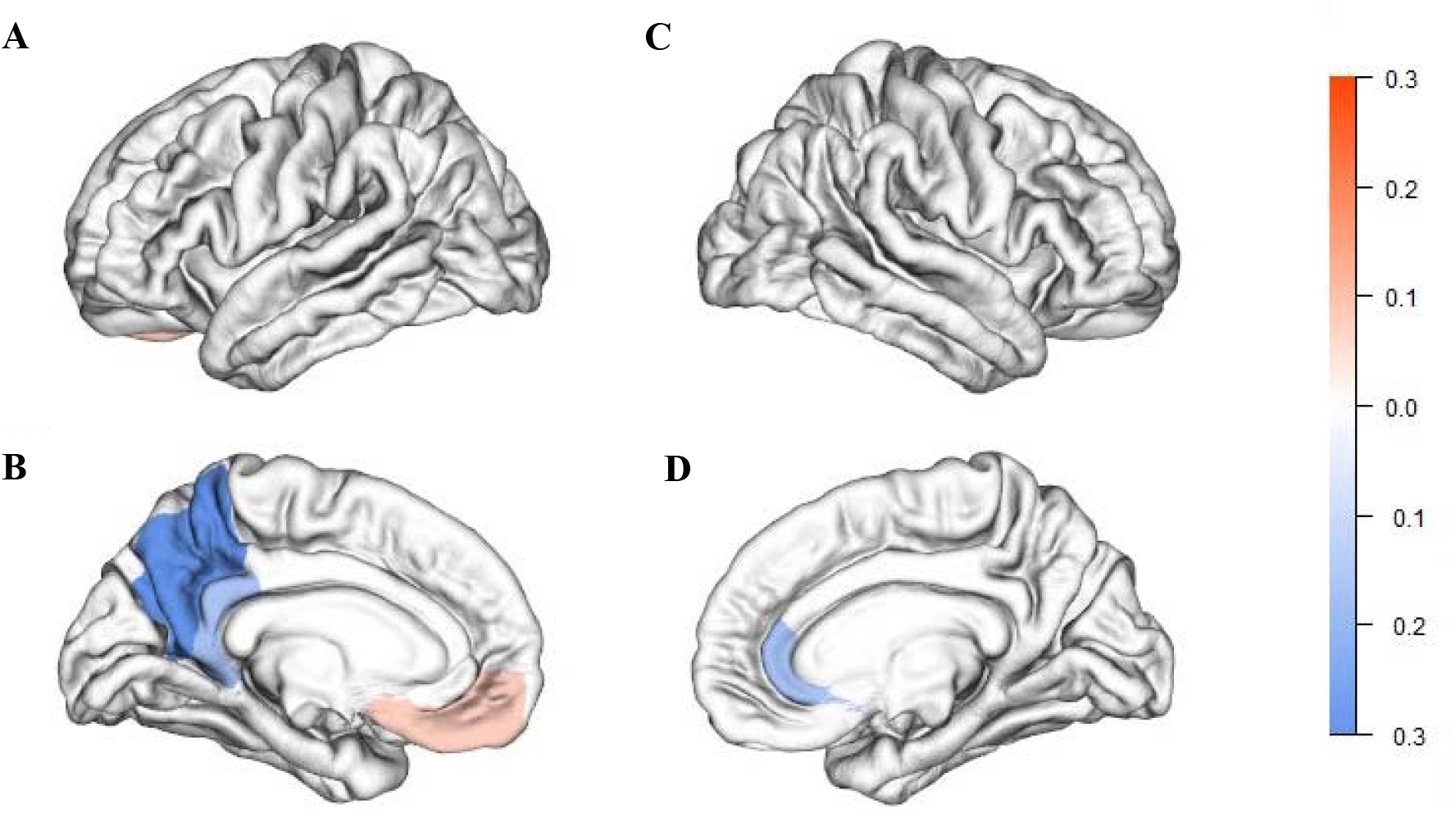
Surface-based visualization of brain regions whose cortical thinning was associated with vmHRV in Cohort 1. A: Left hemisphere lateral view. B: Left hemisphere medial view. C: Right hemisphere lateral view. D: Right hemisphere medial view. Regions shown correspond to predictors retained in the final regression model (Table 1). Color hue indicates the direction of the association (blue: greater cortical thinning associated with higher vmHRV; red: greater cortical thinning associated with lower vmHRV). while color intensity reflects the relative importance of each region for explaining variance in vmHRV.

**Table 1.** Results of the linear regression model identifying brain regions whose cortical thinning predicted HRV in Cohort 1.

| Predictor | Estimate | SE | 95% Confidence<br>Intervall |  | t-value | p-value |
| --- | --- | --- | --- | --- | --- | --- |
|  |  |  | lower | upper |  |  |
| (Intercept) | 55.370 | 5.352 | 44.025 | 66.715 | 10.346 | < 0.001 |
| Left precuneus | -24.177 | 5.528 | -35.896 | -12.458 | -4.373 | <0.001 |
| Left Isthmus cingulate | -13.203 | 5.825 | -25.551 | -0.854 | -2.267 | <0.05 |
| Left medial orbitfrontal | 12.884 | 5.634 | 0.939 | 24.828 | 2.287 | <0.05 |
| Right rostral anteriocingulate | -18.126 | 5.965 | -30.772 | -5.481 | -3.039 | <0.01 |
| $R^2 / R^2$ adjusted | | | | | | 0.72 / 0.65 |

### Cohort 2: development from 12 to 14 years

In the younger cohort (cohort 2), vmHRV was significantly associated with cortical thinning across a distributed set of frontal, parietal and cingulate midline structures as well as temporal regions (see *Figure 2*). Greater cortical thinning in the right pars opercularis, *β* = −10.63, *SE* = 3.53, *t* = −3.012, *p* = .008, and the right inferior temporal cortex, *β* = −17.44, *SE* = 3.78, *t* = −4.61, *p* < .001, was associated with higher vmHRV. In contrast, greater cortical thinning in the right supramarignal gyrus, *β* = 7.38, *SE* = 3.23, *t* = 2.29, *p* = .03, was associated with lower vmHRV. In medial frontal and cingulate regions, cortical thinning showed region-specific associations. Greater thinning in the left isthmus of the cingulate cortex, *β* = 15.86, *SE* = 3.64, *t* = 4.36, *p* < .001, was associated with lower vmHRV, whereas greater thinning in the left medial orbitofrontal cortex was associated with higher vmHRV, *β* = −7.81, *SE* = 3.34, *t* = −2.34, *p* = .03. Another significant link was observed for the left paracentral cortex, *β* = −12.64, *SE* = 3.91, *t* = −3.23, *p* = .005, showing a positive association between cortical thinning and vmHRV. The final model explained 81% of the observed variance in vmHRV (see *Table 2*). Sensitivity analyses demonstrated that the principal cortical regions identified by the stability-selection procedure were largely robust across a broad range of elastic net mixing parameters (α = 0.1–0.9), supporting the stability of the observed regional pattern (Supplementary Table 7). Including additional covariates did not reveal significant effects of age at HRV assessment, sex, or global brain volume change over time (all *p* > .24). Moreover, the extended model did not improve model fit compared to the base model, *F*(9, 4) = 0.71, *p* = .69, indicating that these factors did not meaningfully contribute to explaining vmHRV beyond regional cortical thinning. For detailed results, see Supplementary Table S5.

**Figure 2.**
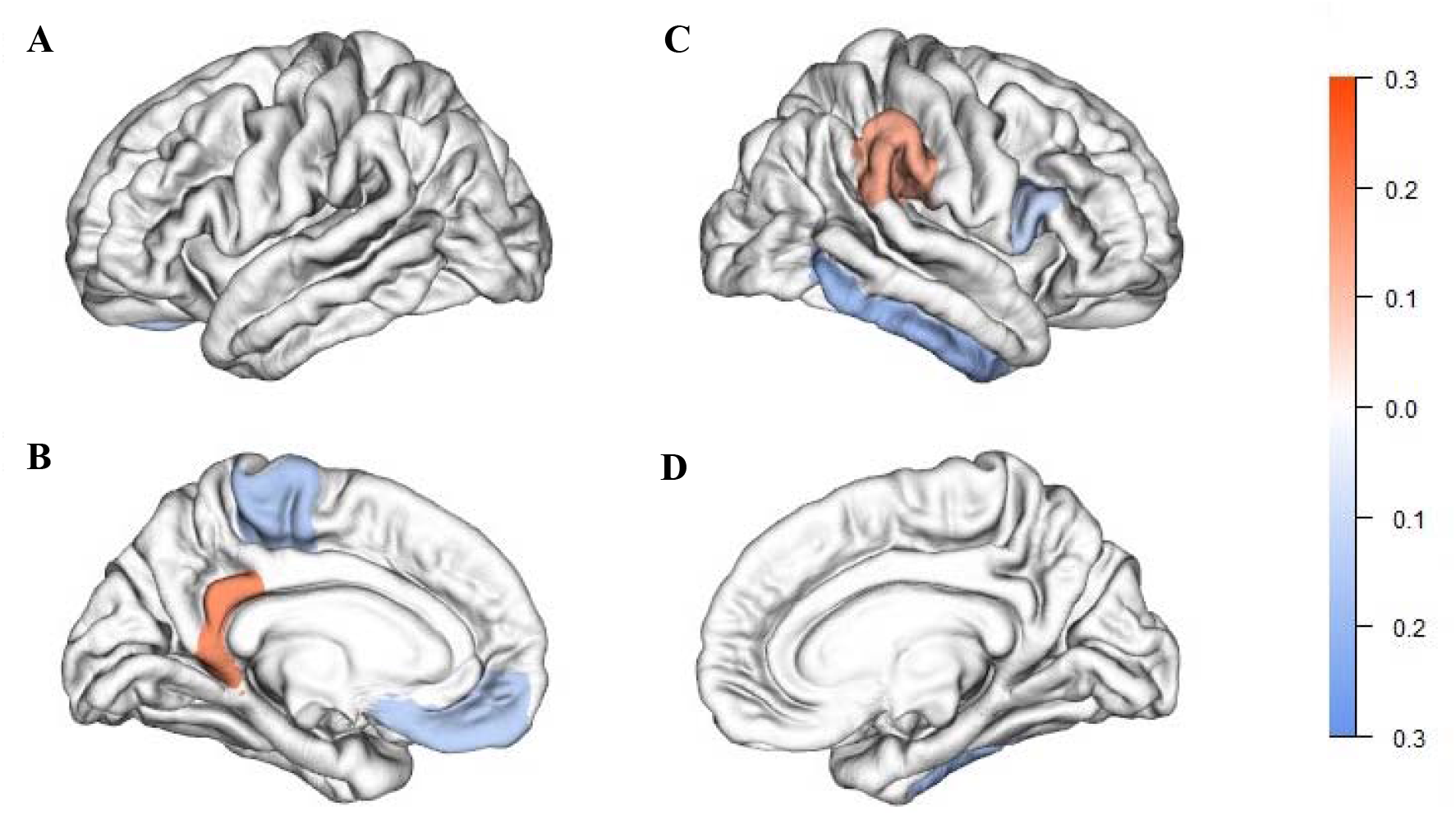
Surface-based visualization of brain regions whose cortical thinning was associated with vmHRV in Cohort 2. A: Left hemisphere lateral view. B: Left hemisphere medial view. C: Right hemisphere lateral view. D: Right hemisphere medial view. Regions shown correspond to predictors retained in the final regression model (Table 2). Color hue indicates the direction of the association (blue: greater cortical thinning associated with higher vmHRV; red: greater cortical thinning associated with lower vmHRV). while color intensity reflects the relative importance of each region for explaining variance in vmHRV.

**Table 2.** Results of the linear regression model identifying brain regions whose cortical thinning predicted HRV in Cohort 2.

| Predictor | Estimate | SE | 95% Confidence<br>Intervall |  | t-value | p-value |
| --- | --- | --- | --- | --- | --- | --- |
|  |  |  | lower | upper |  |  |
| (Intercept) | 54.541 | 2.616 | 48.995 | 60.087 | 20.847 | <0.001 |
| Left medial orbitofrontal | -7.817 | 3.344 | -14.906 | -0.723 | -2.338 | <0.05 |
| Right supramarginal | 7.378 | 3.226 | 0.541 | 14.216 | 2.287 | <0.05 |
| Right pars opercularis | -10.629 | 3.528 | -18.109 | -3.149 | -3.012 | <0.01 |
| Left isthmus cingulate | 15.859 | 3.635 | 8.151 | 23.566 | 4.362 | <0.001 |
| Right inferiotemporal | -17.438 | 3.782 | -25.456 | -9.419 | -4.610 | <0.001 |
| Left paracentral | -12.637 | 3.909 | -20.922 | -4.351 | -3.233 | <0.01 |
| $R^2 / R^2$ adjusted | | | | | | 0.86 / 0.81 |

## Discussion

This study examined whether individual differences in cortical thinning (CT) trajectories are associated with vmHRV in two independent cohorts of children and adolescents. Structural MRI data were acquired using an accelerated longitudinal design at three time points, while vmHRV was measured one year later at follow-up.

In both cohorts, stronger correlations between vmHRV and cortical thinning were observed across distributed cortical regions rather than being confined to a single anatomical structure. Notably, regions showing robust associations included medial and posterior midline structures such as the precuneus and isthmus of the cingulate cortex, as well as orbitofrontal areas. Some of the involved areas, such as the medial prefrontal cortex, the posterior cingulate cortex and adjacent precuneus are part of the default mode network, which is a major neuronal network (Gonen et al., 2020) that supports emotional processing, self-referential mental activity, and autobiographical recollection (Andrews-Hanna, 2009). Beyond these established cognitive and affective functions, contemporary accounts increasingly conceptualize the default mode network as embedded within a broader allostatic system that supports the prediction and regulation of internal bodily or metabolic states in relation to environmental demands (Theriault et al., 2025). Within this framework, processes such as self-referential thinking and autobiographical simulation are not viewed as purely “mental,” but as components of a predictive architecture that helps anticipate bodily needs and guide adaptive regulation. In line with this perspective and our results, developmental changes in default mode network architecture have been linked to the degree of environmental challenge or allostatic load experienced during childhood, suggesting that its structural refinement may, in part, reflect adaptation to cumulative regulatory demands (Rebello et al., 2018). Against this background, the observed associations between vmHRV and cortical thinning in default mode regions may index the coordinated maturation of neural systems supporting both higher-order predictive processing and autonomic regulation.

From a biological developmental perspective, the observed correlations between vmHRV and cortical thinning as a marker of cortical maturation can be interpreted on the basis of three possible, non-mutually exclusive hypotheses, none of which can be directly tested with the present design. First, higher vagal activity may have positive effects on normative cortical thinning during adolescence. This is underscored by studies linking lower vagal activity to increased stress and greater stress susceptibility in children and adolescents (Michels et al., 2013; Porges, 1992). Second, maturational changes reflected by cortical thinning, particularly in regions involved in autonomic regulation such as the cingulate and prefrontal cortex, may influence the developmental trajectory of vagal activity itself. This interpretation is consistent with the findings already discussed on the characteristic developmental course of vagal activity (Silvetti et al., 2001), which suggest that cortical thinning during adolescence may contribute to normative changes in vagal regulation. Third, trajectories of cortical thinning and vagal activity may proceed in parallel independently of each other, or both may be influenced by third factors that were not investigated in the present study.

While specific regional patterns differed between cohorts, there was substantial convergence at the functional network level rather than at the level of individual regions. In both samples, associations involved the isthmus cingulate cortex as well as orbitofrontal and parietal regions, suggesting that vmHRV during adolescence relates to network-level trajectories of cortical thinning within distributed cortical systems. Importantly, the directionality was mixed within this broader network pattern: in several regions (e.g., precuneus & cingulate cortices in Cohort 1), greater cortical thinning predicted higher vmHRV, while in others (e.g., left medial OFC in Cohort 1, left isthmus of cingulate in Cohort 2), greater thinning predicted lower vmHRV.

This pattern of mixed directions contrasts with the findings in adults, where greater cortical thickness is consistently associated with higher vmHRV (Winkelmann et al., 2017; Yoo et al., 2018), and highlights a developmentally driven shift in the relationship between CT and vmHRV across the lifespan (Koenig et al., 2018). Importantly, while specific regional relationships varied across cohorts, convergence occurred at the level of broader cortical systems, suggesting that vmHRV during adolescence is related to distributed maturation at the network level rather than to isolated anatomical regions. It should also be noted that the development of the adolescent brain is characterized not only by changes in cortical thickness but also by increasing functional connectivity between brain regions, which supports the progressive integration of individual subsystems such as the default mode network (Fan et al., 2021). Accordingly, several interacting developmental processes appear to shape the associations between the brain and the autonomic nervous system during this phase, which can lead to regionally heterogeneous patterns that differ from the more uniform relationship observed in adults, where greater CT corresponds to higher vmHRV.

Although the current study supplements cross-sectional data on brain maturation and vagal regulation during adolescence, it nevertheless reveals important aspects that would be useful to examine in the future. Notably, the present analyses were exploratory in nature, allowing us to identify specific brain regions in which developmental trajectories of cortical thinning predicted vagal activity at a later developmental stage. The observation of convergent regional patterns across both cohorts may be interpreted as an initial indication of robustness and external validity. However, given the exploratory approach, these findings should be interpreted with caution, as such analyses are inherently susceptible to inflated false-positive rates, sample-specific effects, and overfitting (Wagenmakers et al., 2012). It should also be noted that the one-year interval between the final structural assessment and the vmHRV measurement reflects the timing of data collection in the broader study protocol rather than a hypothesis-driven design choice, which limits conclusions about the temporal dynamics of this association. Consequently, it is essential that future studies seek to replicate these findings using hypothesis-driven designs with clearly defined regions of interest based on the patterns identified here.

Given that adolescence is a critical period for the emergence of psychiatric disorders (Paus et al., 2008), one further consideration would be to replicate the present findings in samples with greater psychological distress (i.e., adolescents with depression), given that vagal activity is reduced in adolescents with depression (Baumeister-Lingens et al., 2023) and cortical thickness is associated with the first onset of depression in adolescence (Foland-Ross et al., 2015). Another consideration could be to extend the present findings neurodevelopmentally by integrating other measures of autonomic nervous system function and markers associated with vagal activity. The present study is limited to the structural resolution of regional CT, an approach that provides information only about the cortical surface and does not capture structural changes in subcortical regions such as the amygdala or hippocampus (Carnevali et al., 2018). Therefore, possible correlations between vmHRV and the development of deeper brain structures could not be investigated. To address these, other structural measures, such as brain regional volume, would be necessary and should be complimented in future studies. Another direction for future research will be to combine structural measurements of cortical maturation with functional neuroimaging markers of brain networks involved in autonomic regulation. Such multimodal approaches could help clarify whether and how developmental changes in CT are reflected in functional brain networks that support vagal autonomic regulation during adolescence (e.g. Thayer et al., 2012).

In conclusion, the present longitudinal study provides first evidence that individual differences in cortical maturation during adolescence are related to vmHRV. In two independent cohorts, greater cortical thinning in various cortical regions, including cingulate, orbitofrontal, parietal, and medial cortical areas, was associated with higher vmHRV. These findings extend previous cross-sectional studies in adults by suggesting that the association between cortical structure and vagal autonomic regulation is observable during adolescent brain maturation, although the longitudinal, observational nature of the present data does not allow conclusions about the direction or mechanism of this association. Despite the causal direction of these associations remains unclear, the findings underscore that adolescence is a critical developmental window in which the trajectories of cortical maturation and vagal autonomic regulation appear to be statistically associated. Future longitudinal studies incorporating derived hypotheses-driven approaches and additional structural and functional measurements may help to clarify the mechanisms underlying this developmental coupling.

## Data Availability

The datasets generated and analyzed, along with the code used in this study, are available from the corresponding author upon reasonable request.

