## Supplementary Material for "Cortical Thinning Predicts Resting Vagally Mediated Heart Rate Variability: A Longitudinal Study Across Adolescent Development"

Supplementary Table 1. List of used ROI in alphabetical order.

| 1. | Caudal anterior cingulate cortex |
| --- | --- |
| 2. | Caudal middle frontal gyrus |
| 3. | Cuneus cortex |
| 4. | Entorhinal cortex. |
| 5. | Fusiform gyrus |
| 6. | Inferior parietal lobule |
| 7. | Inferior temporal gyrus |
| 8. | Insular cortex |
| 9. | Isthmus of the cingulate cortex |
| 10. | Lateral occipital cortex |
| 11. | Lateral orbitofrontal cortex |
| 12. | Lingual gyrus |
| 13. | Medial orbitofrontal cortex |
| 14. | Middle temporal gyrus |
| 15. | Paracentral lobule |
| 16. | Parahippocampal gyrus |
| 17. | Pars opercularis of the inferior frontal gyrus |
| 18. | Pars orbitalis of the inferior frontal gyrus. |
| 19. | Pars triangularis of the inferior frontal gyrus |
| 20. | Pericalcarine cortex |
| 21. | Postcentral gyrus |
| 22. | Posterior cingulate cortex |
| 23. | Precentral gyrus |
| 24. | Precuneus cortex |
| 25. | Rostral anterior cingulate cortex |
| 26. | Rostral middle frontal gyrus. |
| 27. | Superior frontal gyrus |
| 28. | Superior parietal lobule |
| 29. | Superior temporal gyrus |
| 30. | Supramarginal gyrus |
| 31. | Transverse temporal gyrus |

| Supplementary Table 2. Descriptive statistics of regional thinning slopes per year by hemisphere in Cohort 1. | | | | | | | | | | | | | | |
| --- | --- | --- | --- | --- | --- | --- | --- | --- | --- | --- | --- | --- | --- | --- |
|  | ***Left hemisphere*** | | | | | | | ***Right hemisphere*** | | | | | | |
| ***Region*** | ***Mean*** | ***SD*** | ***Min*** | ***25%*** | ***50%*** | ***75%*** | ***Max*** | ***Mean*** | ***SD*** | ***Min*** | ***25%*** | ***50%*** | ***75%*** | ***Max*** |
| *caudalanteriorcingulate* | *-46.82* | *18.32* | *-96.59* | *-57.40* | *-45.09* | *-35.61* | *-15.12* | *-1.78* | *25.35* | *-61.94* | *-14.57* | *3.47* | *14.42* | *44.94* |
| *caudalmiddlefrontal* | *-25.00* | *8.13* | *-43.19* | *-29.69* | *-26.72* | *-17.36* | *-10.94* | *-37.69* | *15.29* | *-74.90* | *-46.15* | *-38.83* | *-29.59* | *-10.93* |
| *cuneus* | *-24.07* | *22.33* | *-57.60* | *-43.81* | *-26.71* | *-8.84* | *17.79* | *-12.92* | *10.95* | *-33.04* | *-17.32* | *-12.98* | *-8.08* | *10.80* |
| *entorhinal* | *-13.44* | *8.08* | *-26.45* | *-19.76* | *-11.61* | *-8.86* | *1.58* | *-12.84* | *26.69* | *-60.21* | *-29.77* | *-9.96* | *6.46* | *28.96* |
| *fusiform* | *-22.90* | *6.61* | *-38.82* | *-26.51* | *-22.67* | *-19.00* | *-11.95* | *-23.87* | *8.44* | *-42.60* | *-31.43* | *-22.33* | *-17.33* | *-10.47* |
| *inferiorparietal* | *-36.86* | *22.41* | *-82.00* | *-51.81* | *-35.13* | *-26.97* | *26.09* | *-30.89* | *1.82* | *-34.44* | *-32.06* | *-31.32* | *-29.39* | *-27.05* |
| *inferiortemporal* | *-23.25* | *2.49* | *-27.86* | *-24.98* | *-23.53* | *-21.73* | *-18.50* | *-20.43* | *0.88* | *-22.40* | *-20.87* | *-20.38* | *-19.75* | *-19.10* |
| *isthmuscingulate* | *-21.67* | *5.79* | *-36.28* | *-24.62* | *-21.96* | *-19.18* | *-9.16* | *-16.67* | *17.13* | *-41.55* | *-30.04* | *-16.22* | *-4.38* | *23.79* |
| *lateraloccipital* | *-18.37* | *24.47* | *-51.32* | *-35.07* | *-19.39* | *-13.97* | *55.39* | *-22.07* | *13.77* | *-48.62* | *-31.22* | *-23.38* | *-15.42* | *3.72* |
| *lateralorbitofrontal* | *-29.71* | *0.38* | *-30.55* | *-29.89* | *-29.75* | *-29.45* | *-28.92* | *-27.76* | *19.90* | *-74.06* | *-37.75* | *-23.28* | *-11.69* | *3.95* |
| *lingual* | *-19.51* | *1.14* | *-21.37* | *-20.25* | *-19.27* | *-18.75* | *-17.29* | *-32.95* | *4.21* | *-40.47* | *-35.92* | *-32.56* | *-30.66* | *-23.07* |
| *medialorbitofrontal* | *-17.96* | *2.07* | *-22.15* | *-19.30* | *-17.90* | *-16.91* | *-13.53* | *-16.12* | *31.49* | *-72.71* | *-39.14* | *-1.16* | *2.74* | *26.37* |
| *middletemporal* | *-13.70* | *7.65* | *-23.70* | *-20.84* | *-15.18* | *-8.15* | *2.35* | *-40.54* | *6.58* | *-54.02* | *-43.75* | *-40.13* | *-38.14* | *-27.12* |
| *parahippocampal* | *-10.01* | *26.48* | *-50.45* | *-29.68* | *-13.01* | *9.50* | *57.79* | *-24.73* | *3.32* | *-35.39* | *-25.72* | *-24.77* | *-22.78* | *-19.74* |
| *paracentral* | *-32.95* | *22.06* | *-78.85* | *-46.14* | *-34.04* | *-13.67* | *12.37* | *-31.39* | *7.28* | *-41.64* | *-38.23* | *-30.79* | *-25.62* | *-18.26* |
| *parsopercularis* | *-25.67* | *6.63* | *-41.30* | *-27.50* | *-25.75* | *-21.50* | *-14.01* | *-34.22* | *11.75* | *-59.64* | *-42.01* | *-36.07* | *-24.13* | *-17.00* |
| *parsorbitalis* | *-24.44* | *5.50* | *-33.34* | *-27.62* | *-25.36* | *-20.11* | *-14.65* | *-25.85* | *19.46* | *-61.52* | *-39.10* | *-22.76* | *-12.24* | *3.59* |
| *parstriangularis* | *-35.09* | *1.34* | *-37.34* | *-35.87* | *-35.17* | *-34.44* | *-31.53* | *-41.63* | *1.48* | *-45.63* | *-42.43* | *-41.23* | *-40.82* | *-38.75* |
| *pericalcarine* | *-7.02* | *9.17* | *-24.59* | *-12.74* | *-5.63* | *-2.07* | *12.49* | *-12.31* | *20.53* | *-43.48* | *-28.94* | *-14.51* | *1.39* | *26.50* |
| *postcentral* | *-24.45* | *7.49* | *-34.45* | *-29.05* | *-23.09* | *-21.92* | *-4.87* | *-30.61* | *8.11* | *-46.59* | *-35.82* | *-29.25* | *-23.68* | *-18.69* |
| *posteriorcingulate* | *-30.50* | *11.00* | *-49.53* | *-37.79* | *-29.68* | *-21.27* | *-13.80* | *-22.13* | *12.75* | *-52.60* | *-29.41* | *-22.72* | *-14.48* | *1.23* |
| *precentral* | *-15.45* | *5.23* | *-22.46* | *-20.07* | *-16.43* | *-11.72* | *-3.36* | *-24.37* | *16.96* | *-65.21* | *-37.45* | *-23.80* | *-12.25* | *6.46* |
| *precuneus* | *-29.57* | *6.29* | *-41.25* | *-33.60* | *-28.80* | *-25.84* | *-16.81* | *-20.73* | *7.75* | *-36.30* | *-25.17* | *-20.75* | *-15.65* | *-4.39* |
| *rostralanteriorcingulate* | *-24.70* | *5.21* | *-33.94* | *-28.74* | *-25.11* | *-22.20* | *-13.79* | *-11.61* | *27.71* | *-60.03* | *-27.00* | *-8.83* | *-1.79* | *45.53* |
| *rostralmiddlefrontal* | *-36.45* | *3.05* | *-41.94* | *-38.27* | *-35.94* | *-33.83* | *-32.89* | *-46.20* | *9.20* | *-71.80* | *-50.33* | *-45.43* | *-42.10* | *-29.34* |
| *superiorfrontal* | *-30.33* | *12.24* | *-62.98* | *-37.10* | *-29.43* | *-21.44* | *-8.79* | *-33.50* | *2.99* | *-39.03* | *-35.31* | *-32.41* | *-31.32* | *-29.40* |
| *superiorparietal* | *-22.83* | *13.51* | *-58.26* | *-31.69* | *-19.13* | *-14.42* | *-0.03* | *-30.22* | *0.51* | *-31.03* | *-30.53* | *-30.30* | *-29.83* | *-29.32* |
| *superiortemporal* | *-14.48* | *12.81* | *-43.53* | *-21.04* | *-17.58* | *-2.02* | *4.86* | *-21.35* | *6.99* | *-37.06* | *-25.80* | *-21.47* | *-15.93* | *-10.80* |
| *supramarginal* | *-26.17* | *3.32* | *-33.13* | *-27.57* | *-26.67* | *-23.98* | *-19.00* | *-39.55* | *7.68* | *-54.87* | *-43.82* | *-39.49* | *-33.36* | *-28.31* |
| *transversetemporal* | *-11.91* | *13.47* | *-39.68* | *-14.87* | *-10.96* | *1.06* | *8.89* | *-52.51* | *16.17* | *-90.36* | *-57.82* | *-52.24* | *-42.68* | *-21.28* |
| *insula* | *-22.59* | *5.02* | *-34.75* | *-25.66* | *-22.04* | *-18.84* | *-14.98* | *-10.03* | *22.13* | *-61.03* | *-20.25* | *-10.61* | *-0.15* | *36.81* |

| Supplementary Table 3. Descriptive statistics of regional thinning slopes per year by hemisphere in Cohort 2. | | | | | | | | | | | | | | |
| --- | --- | --- | --- | --- | --- | --- | --- | --- | --- | --- | --- | --- | --- | --- |
|  | ***Left hemisphere*** | | | | | | | ***Right hemisphere*** | | | | | | |
| ***Region*** | ***Mean*** | ***SD*** | ***Min*** | ***25%*** | ***50%*** | ***75%*** | ***Max*** | ***Mean*** | ***SD*** | ***Min*** | ***25%*** | ***50%*** | ***75%*** | ***Max*** |
| *caudalanteriorcingulate* | -17.75 | 7.76 | -33.64 | -21.21 | -17.70 | -13.85 | -1.78 | -34.57 | 19.99 | -77.47 | -49.63 | -29.98 | -20.40 | -4.13 |
| *caudalmiddlefrontal* | -13.53 | 14.87 | -36.45 | -26.21 | -16.38 | 0.61 | 12.14 | -1.11 | 7.75 | -19.45 | -5.15 | -1.83 | 2.92 | 21.31 |
| *cuneus* | -16.44 | 2.40 | -23.30 | -17.67 | -15.91 | -15.09 | -12.18 | -19.89 | 6.82 | -36.54 | -21.95 | -20.09 | -16.49 | -6.41 |
| *entorhinal* | -9.63 | 36.53 | -87.54 | -37.03 | -3.05 | 16.79 | 57.37 | -51.95 | 99.11 | -290.15 | -92.56 | -63.70 | 30.82 | 140.52 |
| *fusiform* | -24.98 | 5.85 | -34.18 | -28.85 | -25.29 | -21.03 | -13.11 | -27.95 | 4.33 | -35.13 | -31.21 | -27.63 | -24.54 | -20.65 |
| *inferiorparietal* | -21.69 | 5.39 | -33.10 | -24.58 | -21.71 | -19.00 | -8.56 | -27.20 | 7.93 | -41.10 | -32.78 | -29.44 | -21.26 | -14.83 |
| *inferiortemporal* | -24.44 | 15.96 | -73.26 | -32.32 | -24.31 | -15.44 | 6.28 | -32.05 | 16.40 | -63.67 | -42.07 | -33.51 | -20.32 | -5.42 |
| *isthmuscingulate* | -5.63 | 10.07 | -24.64 | -11.47 | -3.08 | 2.71 | 7.30 | -26.24 | 1.07 | -28.40 | -27.11 | -26.17 | -25.62 | -24.01 |
| *lateraloccipital* | -20.54 | 0.67 | -22.75 | -20.83 | -20.52 | -20.08 | -19.62 | -27.20 | 4.83 | -41.91 | -28.32 | -25.82 | -24.89 | -20.18 |
| *lateralorbitofrontal* | -26.57 | 12.28 | -48.34 | -36.85 | -23.21 | -19.23 | -0.43 | -37.74 | 9.13 | -50.21 | -46.50 | -37.76 | -30.82 | -18.16 |
| *lingual* | -40.00 | 0.41 | -41.15 | -40.22 | -39.93 | -39.78 | -39.37 | -39.35 | 8.75 | -52.61 | -47.15 | -38.77 | -31.88 | -24.65 |
| *medialorbitofrontal* | -22.64 | 9.88 | -43.99 | -28.50 | -24.61 | -14.65 | -6.68 | -61.51 | 34.09 | -167.34 | -80.14 | -54.69 | -39.38 | -5.53 |
| *middletemporal* | -27.76 | 16.65 | -51.65 | -41.37 | -31.08 | -18.28 | 8.49 | -29.08 | 8.32 | -46.48 | -33.66 | -30.91 | -23.05 | -13.74 |
| *parahippocampal* | -10.13 | 20.74 | -54.14 | -25.68 | -11.50 | 0.80 | 42.19 | -20.55 | 3.72 | -26.23 | -23.23 | -20.98 | -18.54 | -11.00 |
| *paracentral* | -4.75 | 16.27 | -27.94 | -16.21 | -9.04 | 6.12 | 29.09 | 15.40 | 20.57 | -12.79 | -0.27 | 15.40 | 20.53 | 66.82 |
| *parsopercularis* | -0.02 | 5.76 | -12.90 | -3.09 | 0.95 | 4.13 | 8.35 | -19.83 | 1.77 | -22.87 | -21.14 | -20.19 | -18.68 | -16.43 |
| *parsorbitalis* | -14.14 | 0.77 | -15.69 | -14.66 | -14.20 | -13.52 | -12.86 | -27.84 | 12.01 | -46.32 | -38.15 | -28.63 | -17.94 | -3.23 |
| *parstriangularis* | -26.81 | 10.80 | -42.33 | -36.30 | -29.29 | -19.20 | -2.39 | -20.25 | 16.84 | -52.02 | -32.62 | -19.20 | -8.08 | 14.73 |
| *pericalcarine* | -19.72 | 11.20 | -41.30 | -29.08 | -19.99 | -10.09 | -0.05 | -26.64 | 6.65 | -37.45 | -32.24 | -29.49 | -21.68 | -13.62 |
| *postcentral* | -15.36 | 26.04 | -86.28 | -31.99 | -5.60 | -2.61 | 27.01 | -31.64 | 34.85 | -131.38 | -41.68 | -23.24 | -7.55 | 11.74 |
| *posteriorcingulate* | -28.02 | 7.49 | -37.17 | -34.78 | -28.86 | -23.81 | -9.04 | -5.59 | 8.66 | -22.35 | -11.19 | -5.92 | -2.14 | 13.98 |
| *precentral* | 10.44 | 18.05 | -57.21 | 6.38 | 12.13 | 21.02 | 32.40 | 28.20 | 11.77 | 6.60 | 20.74 | 29.89 | 34.02 | 56.19 |
| *precuneus* | -24.44 | 3.77 | -30.70 | -27.29 | -24.18 | -22.19 | -16.59 | -21.18 | 9.84 | -45.32 | -25.29 | -21.41 | -17.11 | 4.08 |
| *rostralanteriorcingulate* | -38.92 | 27.42 | -92.95 | -57.76 | -35.51 | -16.83 | 8.90 | -40.86 | 30.53 | -116.43 | -60.91 | -45.71 | -21.26 | 32.87 |
| *rostralmiddlefrontal* | -34.76 | 4.25 | -43.24 | -37.94 | -33.17 | -32.11 | -27.74 | -26.23 | 6.30 | -36.02 | -31.33 | -26.81 | -23.55 | -14.75 |
| *superiorfrontal* | -9.94 | 7.06 | -20.29 | -13.74 | -12.22 | -6.95 | 9.66 | -14.10 | 12.84 | -38.38 | -19.49 | -15.61 | -9.36 | 15.59 |
| *superiorparietal* | -14.65 | 26.02 | -68.53 | -23.87 | -14.04 | 0.16 | 34.02 | -14.93 | 7.18 | -30.86 | -19.63 | -13.68 | -9.61 | -3.96 |
| *superiortemporal* | -17.40 | 9.80 | -50.47 | -21.40 | -16.26 | -12.75 | 2.82 | -22.03 | 9.69 | -54.77 | -25.20 | -22.30 | -16.77 | -3.94 |
| *supramarginal* | -26.73 | 14.40 | -68.55 | -34.08 | -27.50 | -16.23 | -1.09 | -16.68 | 11.73 | -37.03 | -25.11 | -18.24 | -11.20 | 6.86 |
| *transversetemporal* | -12.35 | 5.70 | -21.91 | -16.10 | -13.40 | -8.37 | 0.66 | -29.86 | 6.87 | -44.93 | -34.62 | -27.93 | -25.45 | -15.93 |
| *insula* | -28.80 | 15.37 | -69.94 | -33.63 | -27.64 | -23.43 | 1.59 | -30.81 | 3.10 | -36.04 | -32.18 | -31.31 | -29.20 | -23.91 |

*Supplementary Table 4.* Results of the linear regression model identifying brain regions whose cortical thinning predicted HRV in Cohort 1 including covariates

|  |  |  |  | *95% Confidence Intervall* | |  |  |
| --- | --- | --- | --- | --- | --- | --- | --- |
| *Predictor* |  | *Estimate* | *SE* | *lower* | *upper* | *t-value* | *p-value* |
|  | *Intercept* | *63.579* | *5.770* | *50.527* | *76.631* | *11.020* | *< 0.001* |
| *Left precuneus* |  | *-18.912* | *6.050* | *-32.599* | *-5.225* | *-3.126* | *0.0122* |
| *Left isthmus cingulate* |  | *-12.186* | *7.906* | *-30.071* | *5.698* | *-1.541* | *0.1576* |
| *Left medial orbitofrontal* |  | *16.657* | *5.584* | *4.026* | *29.289* | *2.983* | *0.0154* |
| *Right rostral anteriocingulate* |  | *-13.364* | *6.573* | *-28.233* | *1.505* | *-2.033* | *0.0726* |
| *Sex* |  | *-5.185* | *4.888* | *-16.244* | *5.873* | *-1.061* | *0.3164* |
| *Age (T4)* |  | *-14.833* | *6.541* | *-29.629* | *-0.037* | *-2.268* | *0.0495* |
| *Brain volume slope* |  | *0.003* | *0.001* | *0.0001* | *0.007* | *2.330* | *0.0448* |
| *Left precuneus × sex* |  | *-5.843* | *6.235* | *-19.948* | *8.261* | *-0.937* | *0.3731* |
| *Left isthmus cingulate × sex* |  | *3.718* | *8.592* | *-15.719* | *23.154* | *0.433* | *0.6754* |
| *Left medial orbitofrontal x sex* |  | *-11.553* | *5.617* | *-24.260* | *1.154* | *-2.057* | *0.0699* |
| *Right rostral anteriocingulate x sex* |  | *7.859* | *6.323* | *-6.445* | *22.163* | *1.243* | *0.2453* |
| *R^2^/ R^2^ adjusted* |  |  |  |  |  |  | *0.88 / 0.72* |

*Supplementary Table 5.* Results of the linear regression model identifying brain regions whose cortical thinning predicted HRV in Cohort 2 including covariates

|  |  |  |  | *95% Confidence Intervall* | |  |  |
| --- | --- | --- | --- | --- | --- | --- | --- |
| *Predictor* |  | *Estimate* | *SE* | *lower* | *upper* | *t-value* | *p-value* |
|  | *Intercept* | *73,898* | *23,025* | *19,453* | *128,342* | *3,210* | *0,0149* |
| *Left medial orbitofrontal* |  | *21,325* | *17,387* | *-19,788* | *62,439* | *1,227* | *0,2597* |
| *Right supramarginal* |  | *22,905* | *9,482* | *0,484* | *45,327* | *2,416* | *0,0464* |
| *Right pars opercularis* |  | *-45,048* | *20,941* | *-94,567* | *4,470* | *-2,151* | *0,0685* |
| *Left isthmus cingulate* |  | *4,203* | *12,505* | *-25,366* | *33,772* | *0,336* | *0,7466* |
| *Right inferiortemporal* |  | *-27,545* | *8,855* | *-48,483* | *-6,607* | *-3,111* | *0,0171* |
| *Left paracentral* |  | *3,605* | *16,511* | *-35,438* | *42,648* | *0,218* | *0,8334* |
| *Sex* |  | *-1,508* | *3,468* | *-9,709* | *6,693* | *-0,435* | *0,6768* |
| *Age (T4)* |  | *10,561* | *8,405* | *-9,313* | *30,436* | *1,257* | *0,2492* |
| *Brain volume slope* |  | *-0,002* | *0,002* | *-0,007* | *0,003* | *-0,948* | *0,3747* |
| *Left medial orbitofrontal x sex* |  | *-15,607* | *12,263* | *-44,605* | *13,391* | *-1,273* | *0,2438* |
| *Right supramarginal x sex* |  | *-16,158* | *14,888* | *-51,362* | *19,046* | *-1,085* | *0,3137* |
| *Right pars operculari x sexs* |  | *47,692* | *26,115* | *-14,060* | *109,444* | *1,826* | *0,1106* |
| *Left isthmus cingulate x sex* |  | *11,329* | *10,005* | *-12,329* | *34,988* | *1,132* | *0,2948* |
| *Right inferiortemporal x sex* |  | *18,715* | *9,238* | *-3,129* | *40,560* | *2,026* | *0,0824* |
| *Left paracentral x sex* |  | *-13,999* | *8,022* | *-32,968* | *4,971* | *-1,745* | *0,1245* |
| *R^2^/ R^2^ adjusted* |  |  |  |  |  |  | *0.93 / 0.77* |

| *Supplementary Table 6.* Brain regions showing stable selection across elastic net models (cohort 1). | | | | | | | | | |
| --- | --- | --- | --- | --- | --- | --- | --- | --- | --- |
|  | *Selection rates across different α‘s* | | | | | | | | |
| *Brain region* | *0.1* | *0.2* | *0.3* | *0.4* | *0.5* | *0.6* | *0.7* | *0.8* | *0.9* |
| *Left Precuneus* | *0.62* | *0.58* | *0.57* | *0.52* | *0.52* | *0.47* | *0.47* | *0.45* | *0.43* |
| *Right medialorbitofrontal* | *0.56* | *0.53* | *0.49* | *0.43* | *0.40* | *0.38* | *0.36* | *0.36* | *0.36* |
| *Left isthmuscingulate* | *0.52* | *0.48* | *0.47* | *0.41* | *0.41* | *0.38* | *0.39* | *0.39* | *0.38* |
| *Left medialorbitofrontal* | *0.41* | *0.33* | *0.31* | *0.26* | *0.28* | *0.24* | *0.22* | *0.22* | *0.22* |
| *Right rostral anteriocingulate* | *0.54* | *0.44* | *0.37* | *0.31* | *0.28* | *0.25* | *0.23* | *0.19* | *0.21* |
| *Right isthmuscingulate* | *0.39* | *0.32* | *0.29* | *0.25* | *0.23* | *0.22* | *0.23* | *0.19* | *0.20* |
| *Left precentral* | *0.47* | *0.41* | *0.38* | *0.32* | *0.30* | *0.25* | *0.23* | *0.20* | *0.18* |

| *Supplementary Table 7. Brain regions showing stable selection across elastic net models (cohort 2).* | | | | | | | | | |
| --- | --- | --- | --- | --- | --- | --- | --- | --- | --- |
|  | *Selection rates across different α‘s* | | | | | | | | |
| *Brain region* | *0.1* | *0.2* | *0.3* | *0.4* | *0.5* | *0.6* | *0.7* | *0.8* | *0.9* |
| *Left cuneus* | *0.71* | *0.64* | *0.58* | *0.53* | *0.51* | *0.46* | *0.45* | *0.44* | *0.40* |
| *Right supramarignal* | *0.71* | *0.63* | *0.52* | *0.49* | *0.44* | *0.41* | *0.36* | *0.34* | *0.33* |
| *Left medial orbitofrontal* | *0.62* | *0.55* | *0.50* | *0.47* | *0.46* | *0.43* | *0.40* | *0.40* | *0.36* |
| *Left isthmus cingulate* | *0.65* | *0.58* | *0.50* | *0.45* | *0.42* | *0.37* | *0.37* | *0.33* | *0.31* |
| *Right pars opercularis* | *0.61* | *0.55* | *0.49* | *0.44* | *0.40* | *0.39* | *0.35* | *0.34* | *0.31* |
| *Right fusiform* | *0.66* | *0.57* | *0.50* | *0.43* | *0.38* | *0.34* | *0.32* | *0.30* | *0.27* |
| *Right cuneus* | *0.71* | *0.59* | *0.48* | *0.41* | *0.35* | *0.34* | *0.30* | *0.27* | *0.24* |
| *Right middletemporal* | *0.55* | *0.48* | *0.43* | *0.37* | *0.36* | *0.34* | *0.31* | *0.29* | *0.26* |
| *Right paracentral* | *0.56* | *0.47* | *0.41* | *0.36* | *0.35* | *0.31* | *0.30* | *0.27* | *0.26* |
| *Right inferiortemporal* | *0.60* | *0.47* | *0.38* | *0.34* | *0.30* | *0.27* | *0.23* | *0.21* | *0.20* |
| *Right precentral* | *0.59* | *0.48* | *0.40* | *0.33* | *0.28* | *0.26* | *0.23* | *0.20* | *0.18* |
| *Left paracentral* | *0.50* | *0.40* | *0.33* | *0.27* | *0.27* | *0.24* | *0.22* | *0.19* | *0.21* |
